# Extrapolating Weak Selection in Evolutionary Games

**DOI:** 10.1101/245779

**Authors:** Nanjing U. Zhuoqun Wang, Duke Rick Durrett

## Abstract

This work is inspired by a 2013 paper from Arne Traulsen’s lab at the Max Plank Institute for Evolutionary Biology [10]. They studied the small mutation limit of evolutionary games. It has been shown that for 2×2 games the ranking of the strategies does not change as strength of selection is increased [11]. The point of the 2013 paper is that when there are three or more strategies the ordering can change as selection is increased. Wu et al [10] did numerical computations for fixed *N*. Here, we will instead let the strength of selection *β* = *c/N* and let *N* → ∞ to obtain formulas for the invadability probabilities *ϕ_ij_* that determine the rankings. These formulas, which are integrals on [0, 1], are intractable calculus problems but can be easily evaluated numerically. Here, we concentrate on simple formulas for the ranking order when *c* is small or *c* is large.

## 1 Introduction

Let *a_ij_* be the payoff to a player who uses strategy *i* against an opponent who uses strategy *j*. Let *x_j_* be the frequency of strategy *j* and define the fitness of strategy *i* to be

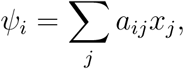

the payoff when a player with strategy *i* plays against a randomly chosen individual. We will study how the frequencies behave under *imitation dynamics*. In this evolution a player *a* chosen at random from the population compares her payoff to an opponent *b* chosen at random and changes her opinion with probability *g*(*β*(*ψ_i(b)_* − *ψ_i(a)_*)) where *β* indicates the strength of selection and *g* : ℝ → (0, 1) is a nondecreasing function, such as the Fermi function *g*(*x*) = 1/(1+*e^−x^*). Thinking of biological competition and following in the footsteps of [4], Wu et al [10] consider the situation in which the mutation rate is small enough so that at most times there is only one strategy *i* in the population. In this case a mutation that introduces one *j* into a population that is all *i* will go to fixation with probability *ϕ_ij_* that can be computed by analyzing a birth and death chain. We suppose that the model rate from *i* to *j μ_ij_* ≡ *μ*. Having computed *ϕ_ij_* one can then analyze the Markov chain with jumps from *i* to *j* at rate *μϕ_ij_* to find the equilibrium frequencies *π_i_*(*β*) of the strategies.

The relative sizes of the *π_i_*(*β*) give an ordering of the strategies. It has been shown that for 2×2 games the ranking of the strategies does not change as strength of fitness is increased [11]. The point of [10] is that when there are three or more strategies the ordering can change as *β* is increased, which the authors tout as a weakness of the “weak selection” viewpoint. In the supplementary materials of [10] they give a complicated argument for the existence of a game in which the strategy ordering can change. In the body of the paper they report on simulations of games with randomly chosen entries that show ranking changes occur in approximately 45% of games.

Here, we will let the strength of selection *β* = *c/N* and let *N* → ∞ to obtain formulas for the invadability probabilities *ϕ_ij_* that determine the rankings. The integrals that give these probabilities, see (6), are difficult to evaluate in general, but they do allow us to compute the ranking order when *c* is small or *c* is large. The dichotomy between small *c* and large *c* is similar to that of *wN* and *Nw* limits introduced by Jeoeng et al [5] and recently studied by Sample and Allen [8]. In the first case one lets the strength of selection *w* → 0 the number of individuals *N* → ∞. In the second the order of the limits is reversed. In our case *w* = *c*?*N* so w → 0 and N → ∞ simultaneously.

The paper is organized as follows: In Section 2 we develop formulas for the *ϕ_ij_*. In Section 3 we compute the stationary distribution for the Markov chain that gives the transition between dominant strategies. In Sections 4 and 5 we give our results for small *c* and large *c* respectively. Section 6 prepares for the study of examples by developing formulas for the three different classes of 2×2 games. In Sections 7 – 11 we apply our results we have obtained to five concrete examples. We summarize our results and discuss open problems in Section 12.

## 2 Formulas for *ϕ_ij_*

In what follows we will use a theorem-proof style of exposition to highlight the main results and to allow the reader to skip the details of the derivations. For simplicity, we will ignore the fact that you can’t play the game against yourself to write the payoffs from playing strategies *i* and *j* when *k* individuals are using strategy *j* as

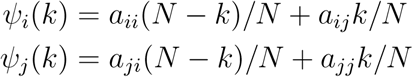

Here, *i* is the wild type, *j* is the mutant. The payoff difference 
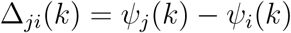
is

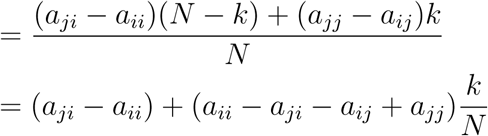

Note that if we add *c_k_* to the entries in the *k*th column, the payoff difference is not changed so we can without loss of generality suppose that the diagonal entries are 0. If we do this then the payoff difference simplifies to

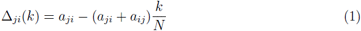

We assume that each individual updates her strategy at rate 1, so the jump rates

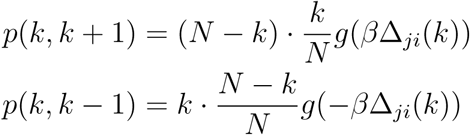

Our first step is to compute the fixation probability. The next result is the same as (4) in the supplementary materials of [10]. In the context of evolutionary games this formula was derived by Taylor et al [9]. However it was first discovered by Karlin and McGregor in the late 1950s [6].

### Theorem 1.

If 
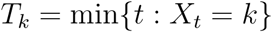
then we have

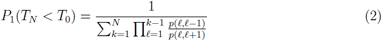

Proof. The first step is to define a function that has *h*(0) = 0, *h*(1) = 1, and 

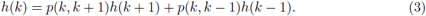

The last equation implies that if we let *X_t_* be the number of the invading type at time *t*, then *E_x_h(X_t_)* remains constant so

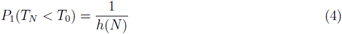

To find *h* we note that (3) implies

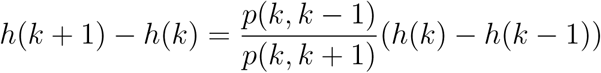

Iterating and using *h*(1) − *h*(0) = 1 it follows that

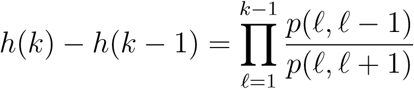

and hence

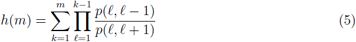

so using (4) gives (2).

### Theorem 2.

If we let *γ* = *g′*(0)/*g′*(0) and suppose *Nβ* → *c* then 
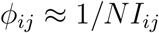
where

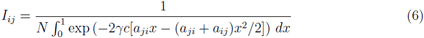

Note that the answer only depends on 2*γc* so we can without loss of generality suppose *γ* = 1/2.

*Proof.* To begin to understand (2) we look at the ratio

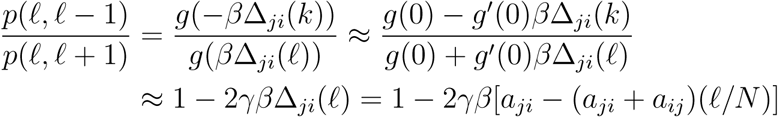

Note that *γ* is the only aspect of g relevant to the value of *ϕ_ij_*. For the Fermi function

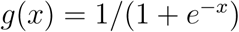
, 
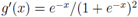
 so *γ* = 1/4.

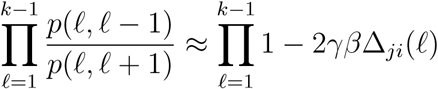

Taking log and using 
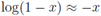
 we have

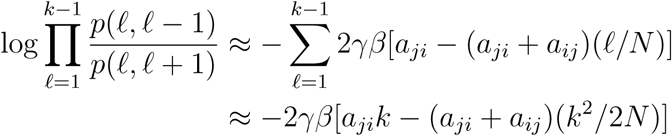

so we have 

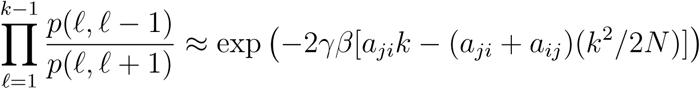

and it follows that if *βN* → *c*

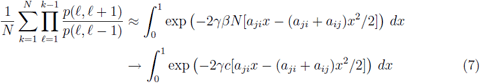

which proves the desired result.

## 3 Equilibrium distribution

If *μ* is the mutation rate from *i* to *j* ≠ *i* the transition rate matrix is

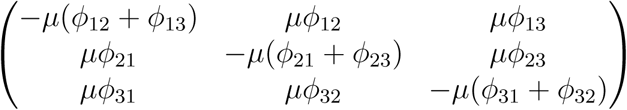

To find the stationary distribution, we need to solve

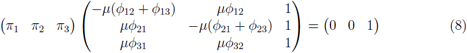

The solution is straightforward, but somewhat tedious, so we just give the answer:

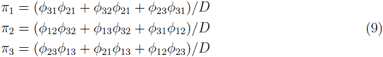

where *D* is the sum of the three numerators. The three formulas can be collapsed into one by writing

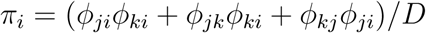

Where *j* = *i* − 1 and *k* = *i* + 1 and the arithmetic is done in ℤ mod 3 = {1, 2, 3}.

## 4 Small *c*

The main result of this section is (11) which allows us to compute the strategy ranking for small *c*. The reader who gets bored with the details can skip to that point. If *c* is small

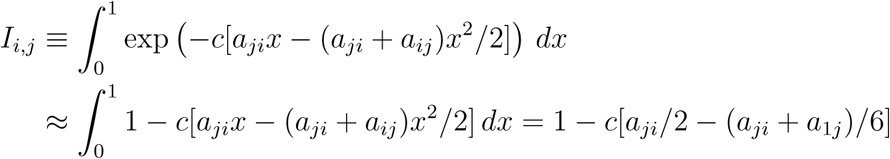

so using (6) 

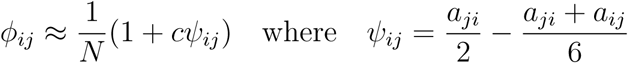

If we (i) cancel the 1/*N*’s and then drop terms of order *c*^2^ then the numerators *n_i_* of the *π_i_* are

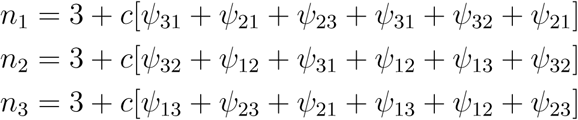

Note that

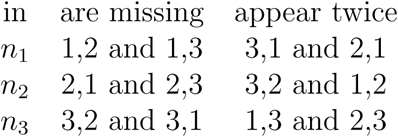

If we let 
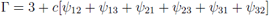
then

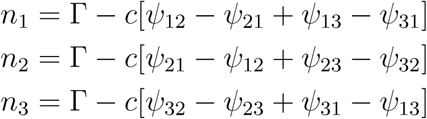

To check the arithmetic, note that the terms that are missing must be positive inside the square brackets while those that appear twice must be negative.

The relative sizes of the *n_i_* gives the strategy ordering for small *c*. To further simplify we note that 
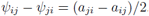
. so if we let

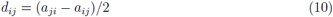

and note that *d_ji_* = −*d_ij_* then we have

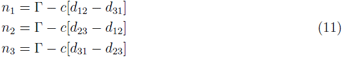

We have used *d*_31_ instead of *d*_13_ so we have the general formula

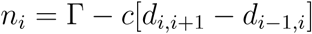

where the arithmetic is done modulo 3. It is not easy to write a formula for the ranking in terms of the entries of the game matrix. However, as the reader will see when we consider examples, it is easy to compute the rankings for a given example.

## 5 Large *c*

The main result of the section is given in the table at the end. The reader who gets bored by the calculus can skip to that point. We are interested in estimating the size of

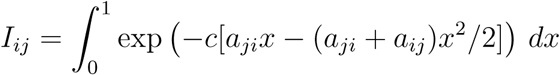

when *c* is large. This will be determined by the largest value of the integrand on [0, 1]. To simplify notation we let *α* = −*a_ji_*, let *β* = *a_ji_* + *a_ij_*, and Artie

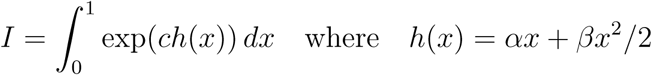

We begin with the case *β* = 0. If *α* < 0

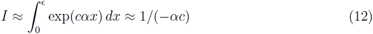

If *α* > 0 we change variables *x* = 1 - *y*

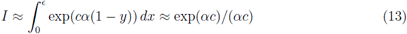

Suppose now *β* ≠ 0. Taking the derivative 
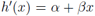
 so *h′*(*x*) = 0 at *x\** = − *α/β*. There are several cases for the location of the maximum of the integrand

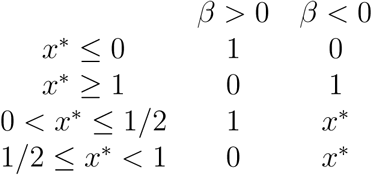

If the maximum occurs at 0 then *h′*(0) = α *<* 0 and

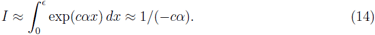

If the maximum occurs at 1 then we change variables *x* = 1 − *y* to get

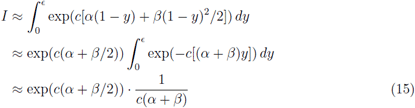

Note that if the maximum occurs at 1 then *α* + *β* > 0. If *β* < 0 then 
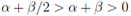
. If β > 0 then we are only in this case when 
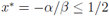
 so *α* + *β*/2 ≥ 0.

If the maximum occurs at 
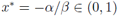
then from the table we see that *β* < 0 and 0 < *α* < −*β*. We change variables *y* = *x* – *x\**

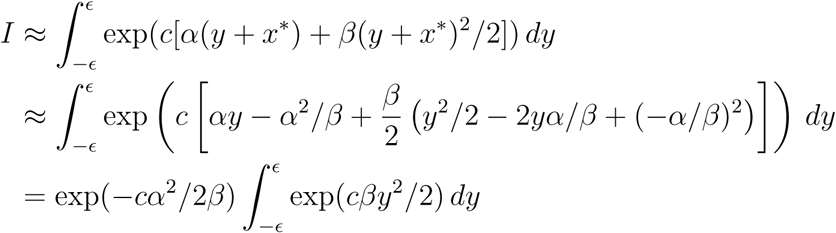

The second and fifth terms in the second line combine to produce the one out front in the third line. The first and fourth cancel. Recalling that normal(0,*σ*^2^) density is

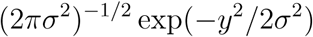

and taking *σ*^2^ = 1/(−*cβ*) we see that

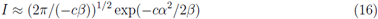

Combining the table that gives the location of the maxima with the last five calculations.

## 6 Three types of edges

To prepare for using the large *c* formulas on examples, we note that *I_ij_* and *I_ji_* only depend on the payoffs in the 2 × 2 subgame with strategies *i* and *j*. In this game only three things can happen *i* ≫ *j*, there is a stable mixed strategy equilibrium, or there is an unstable mixed strategy equilibrium

**Table 1:** Formulas for large *c*

| case | condition | max | $I$ |
| --- | --- | --- | --- |
| I | $\beta > 0, x^* \geq 1/2$ or $\beta < 0, x^* \leq 0$ | 0 | $1/(-c\alpha)$ |
| II | $\beta > 0, x^* \leq 1/2$ or $\beta < 0, x^* \geq 1$ | 1 | $\exp(c(\alpha + \beta/2))/c(\alpha + \beta)$ |
| III | $\beta < 0, 0 < x^* < 1,$ | $x^* = -\alpha/\beta$ | $\exp(-c\alpha^2/2\beta) \cdot (2\pi/(-c\beta))^{1/2}$ |
| IV | $\beta = 0, \alpha < 0$ | 0 | $1/(-c\alpha)$ |
| V | $\beta = 0, \alpha > 0$ | 1 | $\exp(c\alpha)/(c\alpha)$ |

**Case 1.** *i* **dominates** *j*. Since we have 0’s on the diagonal *a_ij_* > 0 and *a_ij_* < 0. We can have *β* = *a_ij_* + *a_ji_* positive or negative.

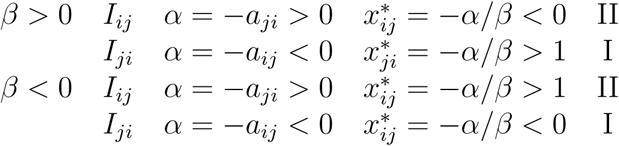

Thus in either case 
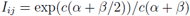
and I_ji_ = 1/(−cα).

**Case 2. Stable mixed strategy equilibrium.** *a_ij_* > 0 and *a_ji_* > 0. Suppose without loss of generality that 
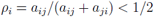

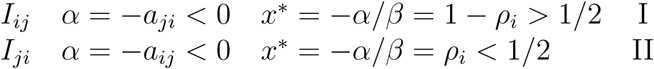

Note that when *i* has *ρ_i_* < 1/2, *I_ji_* is large.

**Case 3. Unstable mixed strategy.** *a_ij_* < 0 and *a_ji_* < 0. In either case

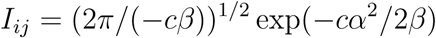

## 7 Example 1. Rock-paper-scissors

The game matrix is

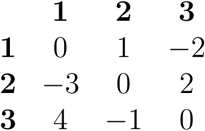

Writing *i* ≫ *j* for strategy i dominates strategy *j* in the (*i*, *j*) subgame this example has 1 ≫ 2 ≫ 3 ≫ 1, which is the same as strategy relationship in the rock-paper-scissors game.

**Figure 1:**
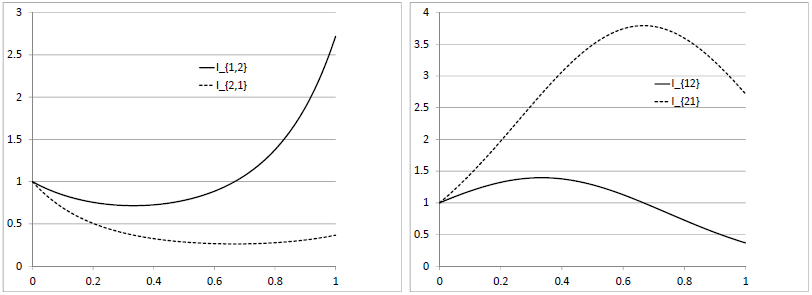
Left: The integrands of *I*_12_ and *I*_21_ when *a*_21_ = 4 and *a*_12_ = 2 (case 2, left) and when *a*_21_ = −4 and *a*_12_ = −2 (case 3, right).

**Small c.** Using (10) and (11)

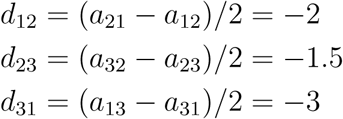

so we have *d*_12_ − *d*_31_ = 1, *d*_23_ − *d*_12_ = 0.5, *d*_31_ − *d*_23_ = −1.5 and it follows that

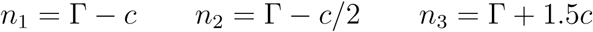

Noting that *n*_3_ > *n*_2_ > *n*_1_, we see that for small *c*, 3 >≻ 2 ≻ 1.

**Large c.** To use the formulas derived in Sections 5 and 6 we compute *α* = −*a_ji_*, *β* = (*a_ji_* + *a_ij_*), and *x\** = −*α*/*β*.

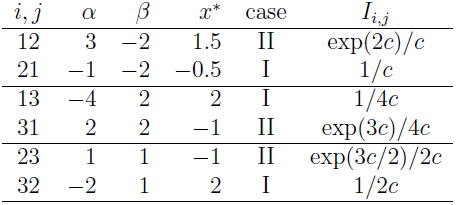

If we recall that (6) implies 
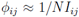
, i.e., a large *I_ij_* means that it is very difficult for *j* to invade *i*, then we see that the answers consistent with the ordering 1 ≫2 ≫3 ≫1: *ϕ_12_*, *ϕ_23_*, and *ϕ_31_* are exponentially small. When we use (9) the *N*’s cancel out. Marking the small terms with *’s

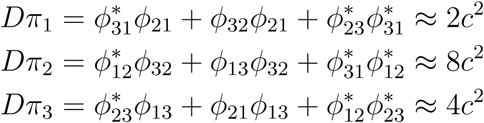

so for large *c* we have 2 ≻ 3 ≻ 1, a change from the small *c* ranking of 3 ≻ 2 ≻ 1.

Generalizing from the concrete example we see that if 1 ≫ 2 ≫ 3 ≫ 1 then all the *d_i,i+1_* are negative. If *d*_31_ is the smallest then

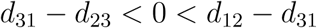

but *d*_23_ − *d*_12_ could be smallest, largest, or in the middle. If *i* ≫ *j* then 
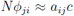
 so we have

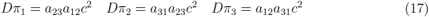

Since the small *c* rankings are based on a linear function of the matrix entries and the large *c* formulas are quadratic, it is not surprising that they can be different.

The asymptotic formula in (17) implies

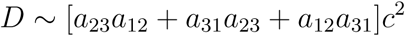

In the concrete example this implies

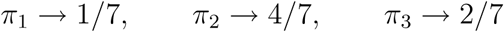

which agrees with Figure 2.

**Figure 2:**
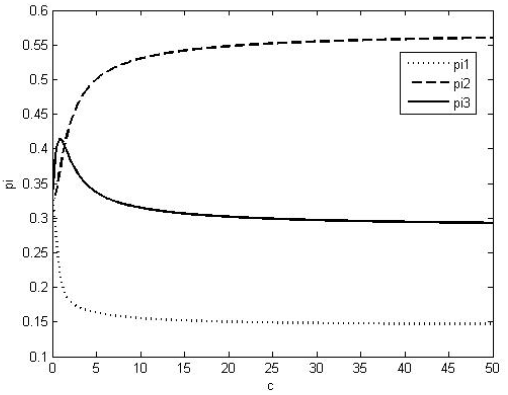
Strategy rankings for Example 1. The ranking changes from 3 ≻ 2 ≻ 1 to 2 ≻ 3 ≻ 1 when 
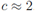
.

## 8 Example 2. One stable edge fixed point

The game matrix is

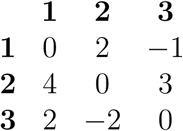

The 1,2 subgame has a mixed strategy equilibrium (1/3, 2/3) in which each strategy has fitness 4/3. When played against this equilibrium 3 has fitness 2/3 − 4/3 = −2/3 so it cannot invade. In the other 2 × 2 subgames 3 ≻ 1 and 2 ≻ 3.

**Small c.** Using (10) and (11)

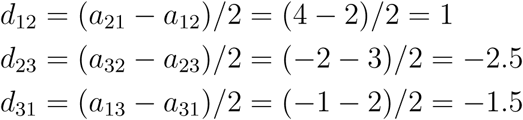

so we have *d*_12_ − *d*_3,1_ = 2.5, *d*_23_ − *d*_12_ = −3.5 and *d*_31_ − *d*_23_ = 1 and it follows that

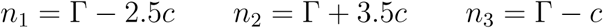

Noting that *n*_2_ > *n*_3_ > n_1_, we see that for small c, 2 ≻ 3 ≻ 1.

**Large c.** Again we need to compute *α* = −*a_ji_*, *β* = (*a_ji_* + *a_ij_*), and *x*^*^ = −*α*/*β*.

| $i, j$ | $\alpha$ | $\beta$ | $x^*$ | case | $I_{ij}$ |
| --- | --- | --- | --- | --- | --- |
| 12 | -4 | 6 | $2/3$ | I | $1/4c$ |
| 21 | -2 | 6 | $1/3$ | II | $\exp(c)/4c$ |
| 13 | -2 | 1 | 2 | I | $1/2c$ |
| 31 | 1 | 1 | -1 | II | $\exp(3c/2)/2c$ |
| 23 | 2 | 1 | -2 | II | $\exp(5c/2)/3c$ |
| 32 | -3 | 1 | 3 | I | $1/3c$ |

Again (6) implies *ϕ*_ij_ = 1/*NI_ij_*, so *ϕ*_21_, *ϕ*_31_ and *ϕ*_23_ are exponentially small. Consulting the formula for the stationary distribution and marking the small terms with ^*^

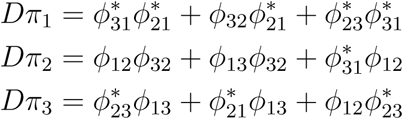

From this we see that 
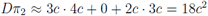
 while π_1_ and π_3_ are exponentially small. To compare π_1_ and π_3_ we need to compute the exact order of the three terms.

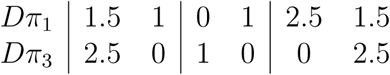

so we have 
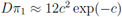
 and 
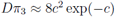
. Thus for large *c* we have 2 ≻ 1 ≻ 3. π_1_ and π_3_ are both exponentially small so π_2_ → 1.

## 9 Example 3. Two stable edge fixed points

The game matrix is

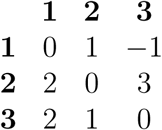

The 1,2 subgame has a mixed strategy equilibrium (1/3, 2/3) in which each strategy has fitness 2/3. When played against this equilibrium, strategy 3 has fitness 2/3 + 2/3 = 4/3 so it can invade. The 2,3 subgame has a mixed strategy equilibrium (3/4, 1/4) in which each strategy has fitness 3/4. When played against this equilibrium, strategy 1 has fitness 3/4 − 1/4 = 1/2 so 1 cannot invade. In the 1,3 subgame 3 ≫ 1

**Small c.** Using (10) and (11)

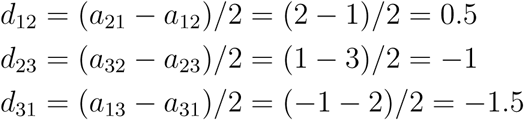

so we have

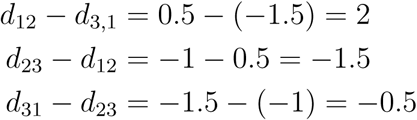

and it follows from (11) that

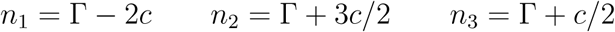

Noting that *n*_2_ > *n*_3_ > *n*_1_, we see that for small *c*, 2 ≻ 3 ≻ 1.

**Large c.** Again, *α* = −*a_ji_*, *β* = (*a_ji_* + *a_ij_*), and *x*^*^ = −*α*/*β*.

| $i, j$ | $\alpha$ | $\beta$ | $x^*$ | case | $I$ |
| --- | --- | --- | --- | --- | --- |
| 12 | -2 | 3 | 2/3 | I | $1/2c$ |
| 21 | -1 | 3 | 1/3 | II | $\exp(c/2)/2c$ |
| 13 | -2 | 1 | 2 | I | $1/2c$ |
| 31 | 1 | 1 | -1 | II | $\exp(3c/2)/2c$ |
| 23 | -1 | 4 | 1/4 | II | $\exp(c)/3c$ |
| 32 | -3 | 4 | 3/4 | I | $1/3c$ |

Using (6), *ϕ_ij_* = 1/*NI_ij_* we see that again *ϕ*_21_, *ϕ*_31_ and *ϕ*_23_ are exponentially small, so marking the small terms with ^*^’s, we have the same pattern as in the previous example.

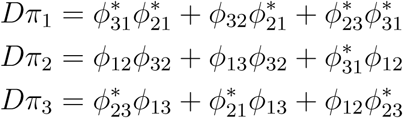

From this we see that

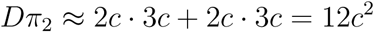

while π_1_ and π_3_ are exponentially small. To compare π_1_ and π_3_ we need to compute the exact order of the three terms.

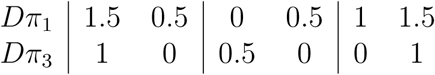

so we have 
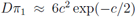
 and 
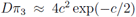
. Thus for large *c* we have 2 ≻ 1 ≻ 3. π_1_ and π_3_ are both exponentially small so π_2_ → 1.

## 10 Example 4. One stable, one unstable

The game matrix is

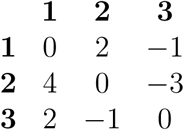

The 1,2 subgame has a mixed strategy equilibrium (1/3, 2/3) in which each strategy has fitness 4/3. When played against this equilibrium 3 has fitness 2/3 − 2/3 = 0 so it cannot invade. The 2,3 subgame has an unstable mixed strategy equilibrium (3/4, 1/4) which can be invaded by 1. In the 1,3 subgame 3 ≫ 1.

**Small c.** Using (10) and (11)

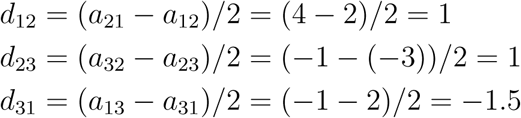

so we have

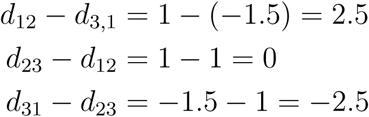

and it follows that

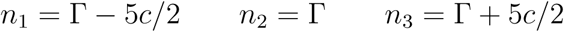

Noting that *n*_3_ > *n*_2_ > *n*_1_, we see that for small *c*, 3 ≻ 2 ≻ 1.

**Large c.** Again, *α* = − *a_ji_*, *β* = (*a_ji_* + *a_ij_*), and *x*^*^ = *α*/*β*. We have an instance of case III here. In this case 
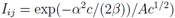
 where 
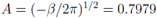

| $i, j$ | $\alpha$ | $\beta$ | $x^*$ | case | $I$ |
| --- | --- | --- | --- | --- | --- |
| 12 | -4 | 6 | 2/3 | I | $1/4c$ |
| 21 | -2 | 6 | 1/3 | II | $\exp(c)/4c$ |
| 13 | -2 | 1 | 2 | I | $1/2c$ |
| 31 | 1 | 1 | -1 | II | $\exp(3c/2)/2c$ |
| 23 | 1 | -4 | 1/4 | III | $\exp(c/8)/A c^{1/2}$ |
| 32 | 3 | -4 | 3/4 | III | $\exp(9c/8)/A c^{1/2}$ |

Using (6), *ϕ*_ij_ = 1/*NI_ij_* we see that again *ϕ*_21_, *ϕ*_31_, *ϕ*_23_, and *ϕ*_3,2_ are exponentially small, so marking the small terms with ^*^’s, we have

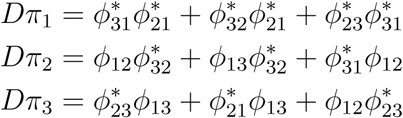

This time all three terms are exponentially small. The exponential orders of the terms are

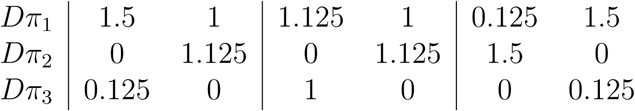

so we have

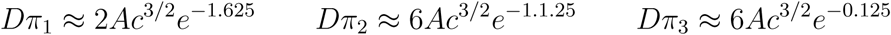

This the ranking remains the same for large *c*: 3 ≻ 2 ≻ 1.

**Figure 5:**
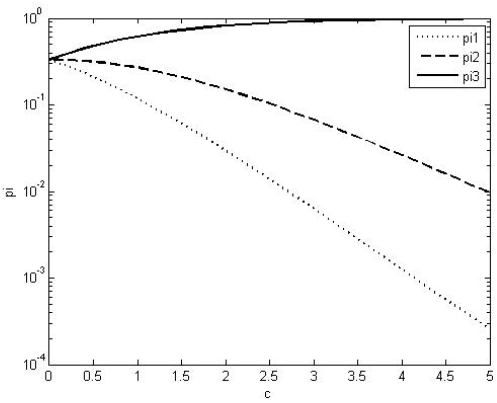
In Example 4 the strategy ranking 3 ≻ 2 ≻ 1 does not change. As *c* → ∞, π_3_ → 1 exponentially fast. In contrast to the two previous examples the exponential decay rates for π_1_ and π_2_ are different.

## 11 Example 5. Two ranking changes

In the previous four examples there has been a most one ranking change. We now give an example with two changes. The game matrix is

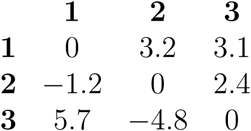

The 1,3 subgame has a mixed strategy equilibrium (3.1/8.8,5.7/8.8) in which each strategy has fitness (5.7)(3.1/8.8) = 2.008. When played against this equilibrium 2 has fitness [(−1.2)(3.1) + (2.4)(5.7)]/8.8 = 1.13, so 2 cannot invade. In the other two 2 × 2 subgames 1 ≫ 2 and 2 ≫ 3.

**Small c.** Using (10) and (11)

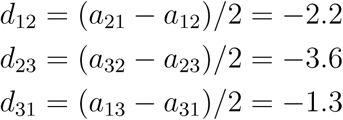

so we have

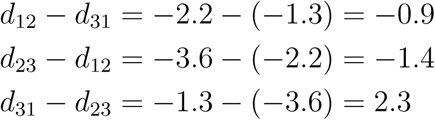

and it follows that

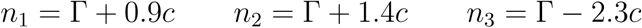

Noting that *n*_2_ > *n*_1_ > *n*_3_, we see that for small *c*, 2 ≻ 1 ≻ 3.

**Large c.** Again, *α* = −*a_ji_*, β = (*a_ji_* + *a_ij_*), and *x*^*^ = −*α*/*β*.

| $i, j$ | $\alpha$ | $\beta$ | $x^*$ | case | $I$ |
| --- | --- | --- | --- | --- | --- |
| 12 | 1.2 | 2 | -0.6 | II | $\exp(2.2c)/(3.2c)$ |
| 21 | -3.2 | 2 | 1.6 | I | $1/3.2c$ |
| 13 | -5.7 | 8.8 | 0.6477 | I | $1/5.7c$ |
| 31 | -3.1 | 8.8 | 0.3523 | II | $\exp(1.3c)/5.7c$ |
| 23 | 4.8 | -2.4 | 2 | II | $\exp(3.6c)/2.4c$ |
| 32 | -2.4 | -2.4 | -1 | I | $1/2.4c$ |

Using (9), *ϕ_ij_* = 1/*NI_ij_* we see that *ϕ*_21_, *ϕ*_31_, and *ϕ*_32_ are exponentially small, so marking the small terms with ^*^’s, we have the same pattern as in the previous example.

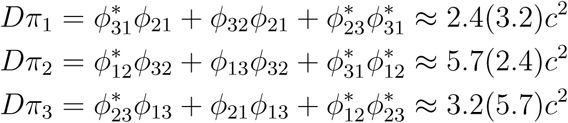

so for large *c* we have 3 ≻ 2 ≻ 1. Comparing this with the small *c* ranking of 2 ≻ 1 ≻ 3 we can see that there must be two ranking changes to bring strategy 3 from last to first.

## 12 Discussion

In this paper we have developed methods for computing strategy rankings in evolutionary games when mutation rates are small, the strength of selection is *c*/*N*, and the population size *N* → ∞. For any *c* the limiting rankings can be computed by numerically evaluating an integrals on [0, 1]. We have simple explicit results when *c* is small or *c* is large that allow us to infer that ranking changes have occurred.

**Figure 6:**
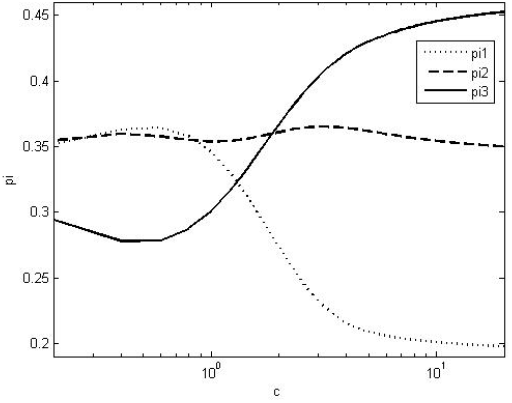
Plot of rankings versus log(*c*) in Example 5.

Simulations of games with 3 ≤ *n* ≤ 9 strategies show (see Figure 4 in [10]) that a positive fraction of *n* strategy games have ≥ *n* − 1 ranking changes, while Figure 3 gives a 3 strategy example that has 4 strategy changes. One can, of course, investigate ranking changes in games numerically, but it would be nice to develop mathematical methods to determine the number of changes for intermediate *c*.

**Figure 3:**
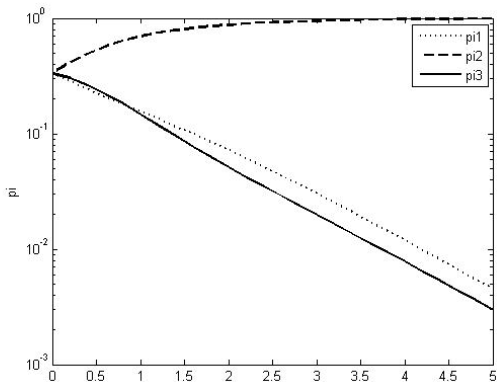
Strategy rankings for Example 2 as a function of *c*. Frequencies are plotted on a log scale so that we can see π_1_ and π_3_ have the same exponential decay rate but π_1_ has a larger constant.

**Figure 4:**
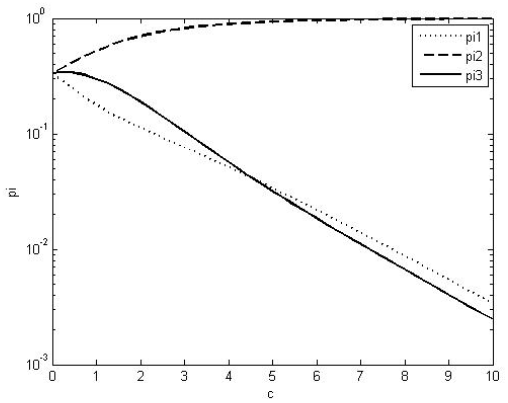
Strategy rankings in Example 3 as a function of *c*. Again the exponential decay rates are the same in π_1_ and π_3_.

The formulas we have for rankings for small *c* and large *c* are simple. They allow us to obtain insights into conditions under which strategy rankings change in rock-paper-scissors examples. However, at this point it does not seem possible to give meaningful results identifying the class of games that have no strategy changes. Perhaps there is no simple answer, but it would be interesting if one could be found.

## Acknowledgments

We would like to thanks Ben Allen and Philipp Altrock for comments on the paper and for bringing some important references to our attention. Arne Traulsen and Bin Wu made many useful suggestions and helped us refocus the paper on our positive contributions.

